# Clinical considerations on antimicrobial resistance potential of complex microbiological samples

**DOI:** 10.1101/2024.05.14.594174

**Authors:** Norbert Solymosi, Adrienn Gréta Tóth, Sára Ágnes Nagy, István Csabai, Csongor Feczkó, Tamás Reibling, Tibor Németh

## Abstract

Antimicrobial resistance (AMR) is one of our greatest public health challenges. Targeted use of antibiotics (AB) can reduce the occurrence and spread of AMR and boost the effectiveness of treatment. This requires knowledge of the antibiotic susceptibility (AS) of the pathogens involved in the disease. Therapeutic recommendations based on classical antibiotic susceptibility testing (AST) are based on the analysis of only a fraction of the bacteria present in the disease process. Next and third generation sequencing technologies allow the identification of antimicrobial resistance genes (ARGs) present in a bacterial community. Using this genomic approach, we can map the antimicrobial resistance potential (AMRP) of a complex, multi-bacterial microbial sample. The same approach can be used to identify antibiotics without any ARGs in the sample that interfere with their activity. Our paper summarises the clinical interpretation opportunities of genomic analysis results from 574 *Escherichia coli* strains and a complex microbiological sample from canine external otitis. In clinical metagenomics, AMRP may be an important approach to make AB therapy more targeted and effective.

**Graphical abstract:** 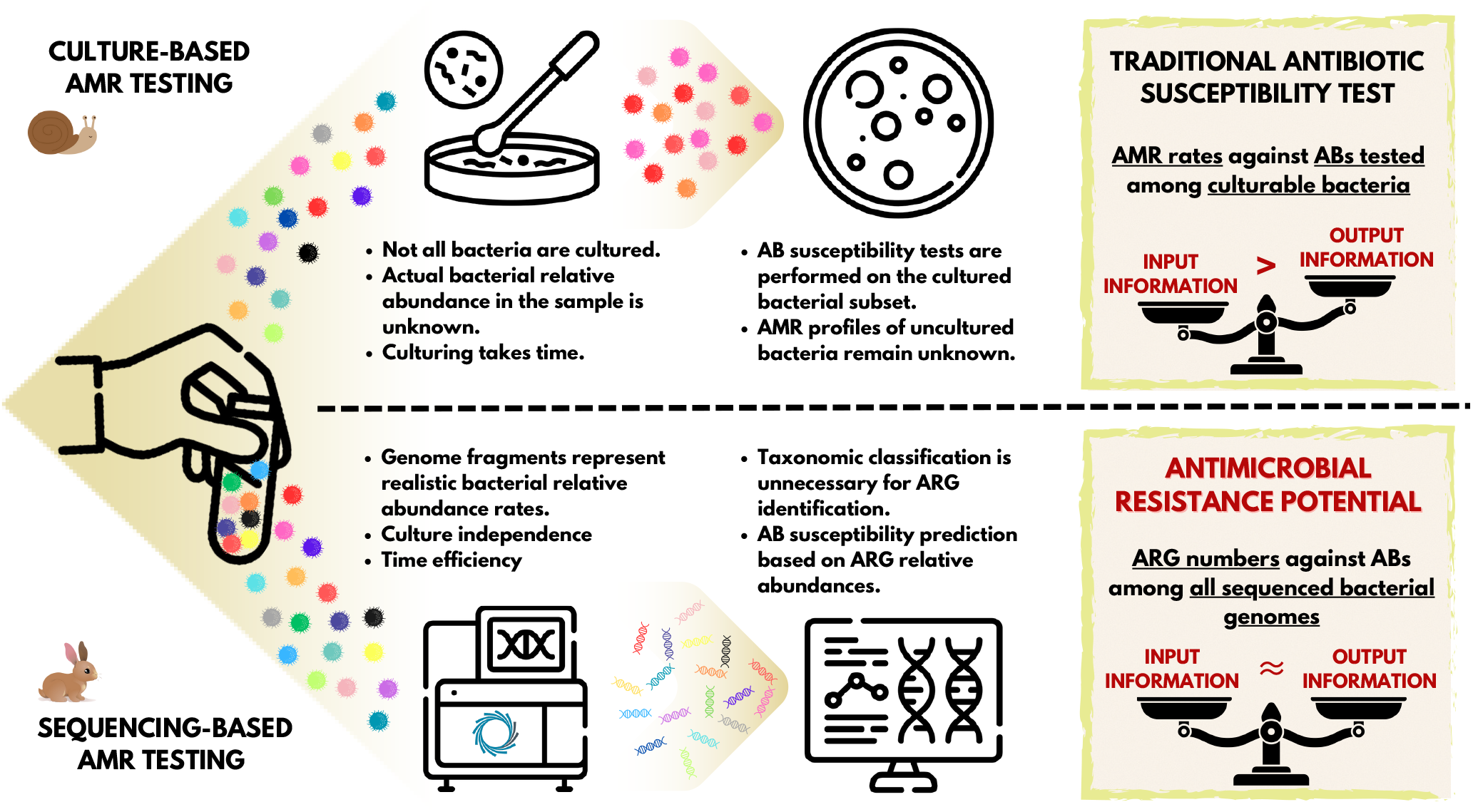

## Introduction

The appearance and use of antibiotics has brought an unprecedented advance in the treatment of bacterial infections.^1^ The general consensus is that the case-specific, targeted use of antibiotics is favorable in clinical settings. This means using antibiotic compounds that are supposed to be effective against the bacteria involved in the disease process to be treated.^2^ The classic standard is to take a sample from the patient for microbiological testing, and perform the bacterial culturing and the antibiotic susceptibility testing (AST) of it.^3^ Based on these tests, we obtain information regarding the set of antibiotics that the bacteria in the culture showed susceptibility to, and the set of compounds to which they are moderately or not at all susceptible.^4^ The general procedure for this widely-used routine bacteriology technique is to first obtain a mixed culture from the sample, then select certain colonies for pure culture,^5^ and perform disk diffusion or microbroth dilution AST.^5^ Importantly, even in a mixed bacterial culture, only a subset of the microorganisms and bacteria involved in the pathological process are represented. This is partly due to the fact that a high proportion of the bacteria ever discovered are unculturable.^6^. Furthermore, even in case of cultivable species, providing the same culture conditions (e.g. type of medium) favors the growth of some taxa over others.^7^ The latter issue can be avoided by creating mixed cultures in parallel, providing different culture environments.^8^ Nevertheless, antibiotic susceptibility testing is always performed on an unpredictably small subset of bacterial strains. Additionally, several phases of traditional microbiological testing are time-consuming processes that take at least a few days to complete. Furthermore, the entire conventional diagnostic workflow is rather labour-intensive.^3^ At the end of the diagnostic process, the phenotypic antimicrobial resistance profile delivered to clinicians refers to a fraction of the bacteria present in the disease process.

At the same time, next-generation sequencing facilitates the identification of a wide variety of genes, including antimicrobial resistance genes (ARGs). Moreover, the identification of the ARGs in microbiological samples can be performed in a considerably shorter time, regardless of whether the bacteria carrying them can be cultured or not, or whether they are considered pathogenic or not.^3^ While the classical approach outlined above yields a phenotypic antibiotic susceptibility profile, ARG detection yields a genotypic antimicrobial resistance profile.^5^ The antibiotic resistance detected in phenotypic settings reflect the activity and functionality of the genetic background^9^, whether or not the genes or polymorphisms that contribute to the generation of the expressed effect, AMR, are known. In the case of a genotypic resistance profile, we do not know whether the gene is expressed or not. However, we can assume that there is a possibility for the expression of the ARGs present that can be interpreted as antimicrobial resistance potential (AMRP).

In our study, the initial step was to investigate the concordance of the phenotypic antibiograms and the ARG content of 574 *Escherichia coli* strains from 5 BioProjects. Based on this result, we describe the AMRP of a complex microbiome and the therapeutic considerations that can be deduced from it.

## Methods

To investigate phenotypic and genotypic antimicrobial resistance, antibiograms and Illumina sequencing data of *E. coli* strains used by Pataki et al.^10^ were downloaded from the European Nucleotide Archive (ENA) at EMBL-EBI repository (https://www.ebi.ac.uk/ena/browser/home). Of these *E. coli* datasets, BioProjects that contained at least 10 samples and an antibiogram involving more than one antibiotic compound were included in the analyses. In the antibiograms, the resistant/intermediate/susceptible categories for each antibiotic were defined by the authors publishing the datasets based on different AST standards and cut-off versions in each BioProject: PRJDB7087 (European Committee on Antimicrobial Susceptibility Testing, EUCAST, 2018), PRJEB14086 (EUCAST, 2013/652/EU), PRJEB21880 (EUCAST, 2015), PRJEB21997 (EUCAST, 2017), PRJNA266657 (Clinical and Laboratory Standards Institute, CLSI, 2015). Raw FASTQ files were obtained from each BioProject. Bioinformatic analyis of the FASTQ files began with quality control, followed by the trimming and filtering of the raw short reads usingTrimGalore (v.0.6.6, https://github.com/FelixKrueger/TrimGalore, accessed on 8/12/2023), setting a quality threshold of 20. More than 50 bp long reads were assembled to contigs by MEGAHIT (v1.2.9)^11^ using default settings.

The metagenomic sample was collected from the right external ear canal of a 4-year-old neutered male Belgian griffon before Total Ear Canal Ablation (TECA) or Lateral Bulla Osteotomy (LBO) surgery. The dog was previously diagnosed with an aural mass submitted as an inflammatory polyp and end-stage external otitis. The detection of the symptoms took place around six weeks before the surgery. Multiple conservative treatment rounds with antibiotic and anti-inflammatory agents had been performed previously with no success. No tumor tissues were identified via histopathology examination. The history of the dog did not include severe allergies. The skin of the ear canal was red, swollen, scaled, thickened, and covered with bloody, purulent ear discharge before the surgery. By the routine bacteriology testing *Pseudomonas aeruginosa* and, in small amounts, *Malassezia pachydermatis* were detected from the sample. Even though, the identified *Pseudomonas aeruginosa* strain appeared to be multi-resistant, ceftazidime, gentamicin, tobramycin, ciprofloxacin, marbofloxacin and polymyxin B were indicated to be potentially effective, if administered locally, in great concentrations. DNA extraction was performed with QIAamp PowerFecal Pro DNA Kit from Qiagen according to the manufacturer’s instructions. The concentrations of the extracted DNA solutions were evaluated with an Invitrogen Qubit 4 Fluorometer using the Qubit dsDNA HS (High Sensitivity) Assay Kit. The metagenomic long-read library was prepared by the Rapid Barcoding Kit 24 V14 (SQK-RBK114.24) from Oxford Nanopore Technologies (ONT). The sequencing was implemented with a MinION Mk1C sequencer using an R10.4.1 flow cell from ONT. The basecalling was performed using dorado (https://github.com/nanoporetech/dorado, v0.4.3) with model dna_r10.4.1_e8.2_400bps_fast@v4.2.0, based on the POD5 files converted from the FAST5 files generated by the sequencer. The raw reads were adapter trimmed and quality-based filtered by Porechop (v0.2.4, https://github.com/rrwick/Porechop) and Nanofilt (v2.6.0)^12^, respectively.

For the Illumina short read samples, the ARG content of the sequence in the contigs and for the ONT metagenome sample in the long reads was identified using ResFinder (v4.5.0) (minimum coverage 60%, minimum identity 90%).^13^

In the agreement analysis between phenotypic AMR and AMR prediction based on ARG detection, we only used AST results that did not use AB combinations, as the genetic background of resistance to single antibiotics is better understood than ARG determination of resistance to AB combinations. From AST results, only resistant and susceptible results were used. Intermediate phenotypic results were excluded as their genotyping was beyond our aim here. Antibiotics negatively affected by the detected ARGs were recorded and collected by each sample.

Based on the AST and ARG results, 2×2 cross-tabulations (TP: true positive, TN: true negative, FP: false positive, FN: false negative) were generated by each antibiotic compound. These were used to calculate metrics to describe the predictive goodness of the genome-based prediction of the phenotype. Phenotypic AST results were the reference (ground truth). Accordingly, the TP group refers to strains with both phenotypic and genotypic resistance to a given antibiotic compound, TN to strains with both phenotypic and genotypic sensitivity, FP to strains with phenotypic sensitivity and genotypic resistance, and FN to strains with phenotypic resistance without the detection of any ARGs to a given drug. In addition to sensitivity (SE=TP/(TP+FN)), specificity (SP=TN/(TN+FP)), negative predictive value (NPV=TP/(TP+FP)) and positive predictive value (PPV=TP/(TP+FP)), point and 95% CI estimates were also calculated for false resistance (major error rate, ME) and false susceptibility (very major rate, VME). As no intermediate category was used, ME = FP / (TN+FP) and VME = FN / (TP+FN).

Internationally accepted guidelines and standards are also available to assess the accuracy of the different AST methods in relation to standard reference methods, i.e. to determine VME and ME values. In the United States, the FDA (Food and Drug Administration) regulations for the marketing of different antibiotic susceptibility testing devices are quite strict. Thus, the ME of the test device examined must not exceed 3% when testing the efficacy of different bacterial species-drug combinations. In case of VME, the proposed statistical criteria include an upper 95% confidence limit for the VME of 7.5% at most, and a lower 95% confidence limit for the VME of 1.5% at most.^14^ An international standard has also been established for the evaluation of antibiotic susceptibility testing devices, which proposes similar but not identical criteria as acceptable limits of accuracy.^15^ The above guidelines and standards have been tailored to compare different phenotypic techniques rather than phenotypic and genotypic methods, which have the potential for greater variation. Thus, we have used the 5% margin of error used in statistics.

## Results

In Figures 1. and 2. we summarize the point and 95% CI estimates of the agreement, prediction goodness metrics for the phenotypic (ground truth) and genotype-predicted AMR, by antibiotics and BioProjects.

**Figure 1.**
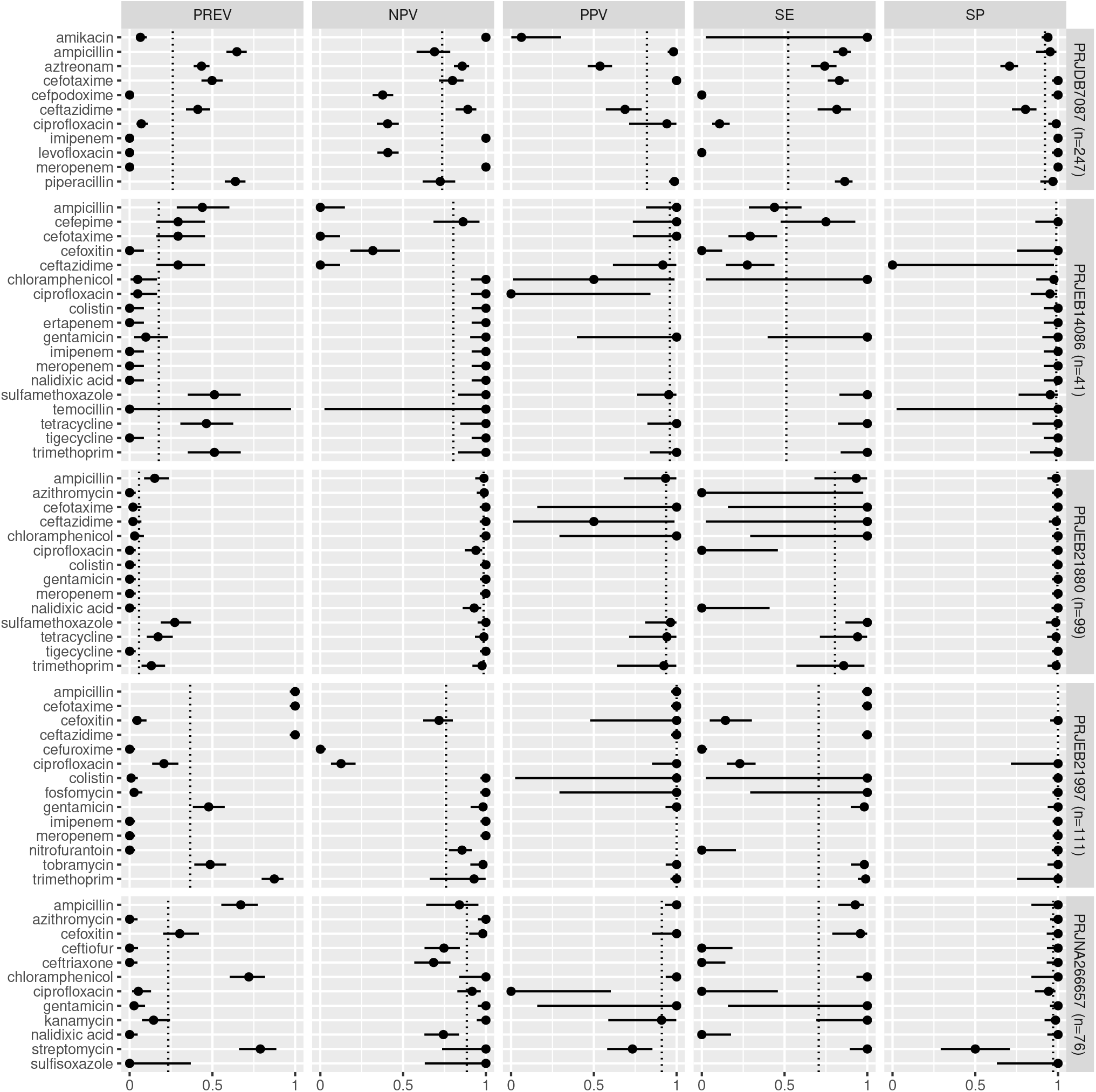
Predictive performance. Antimicrobial resistance gene-based prediction compared to the antibiotic susceptibility test results using five BioProject datasets. Column PREV shows the prevalence (with 95% CI) of phenotypic antimicrobial resistance against certain antibiotics within the BioProjects. Point and 95% CI estimates for negative and positive predictive values, sensitivity, and specificity are presented in columns NPV, PPV, SE, and SP. The dotted vertical lines represent the mean of the metrics within the given BioProject.

**Figure 2.**
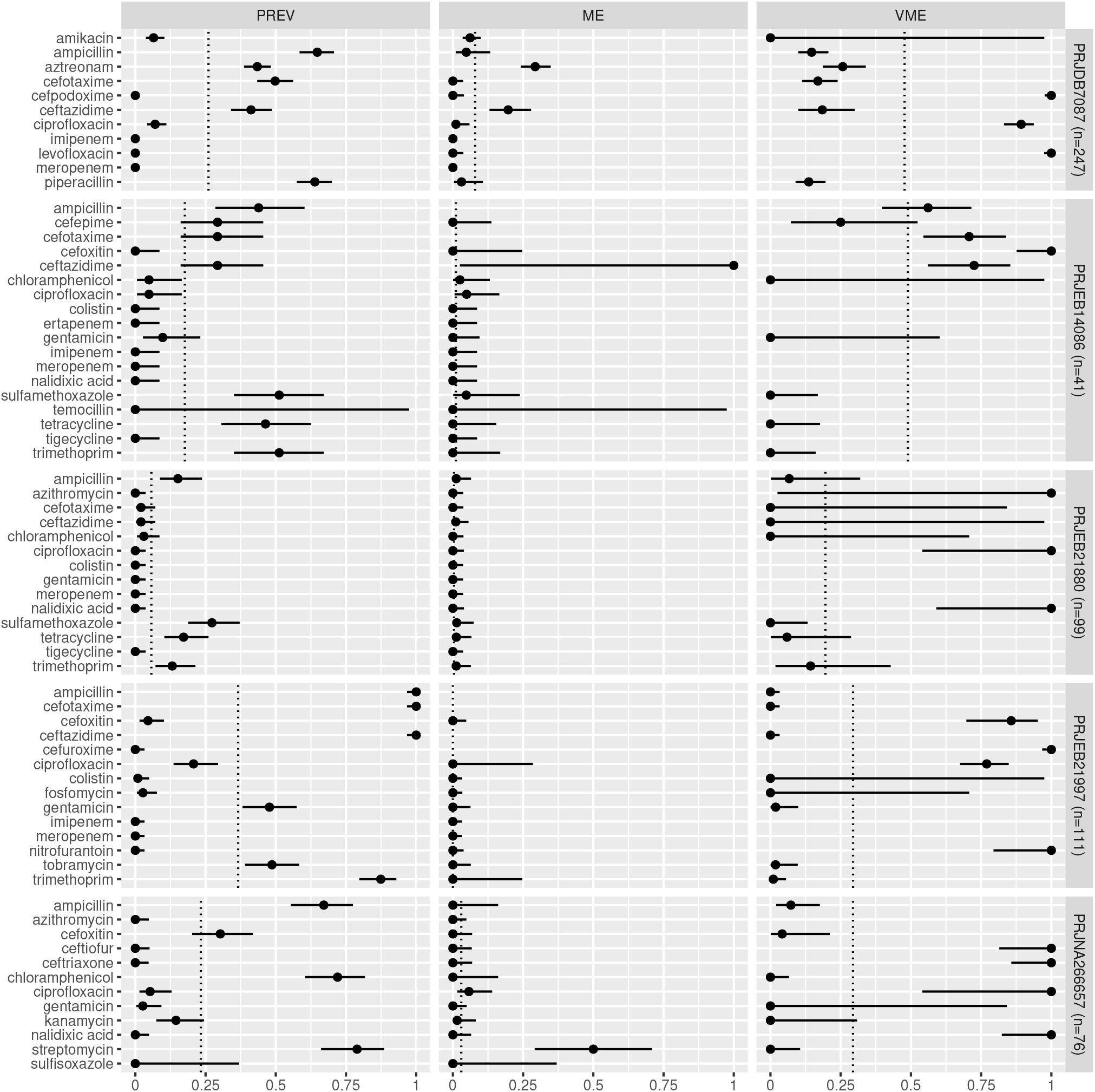
Predictive performance. Antimicrobial resistance gene-based prediction compared to the antibiotic susceptibility test results using five BioProject datasets. Column PREV shows the prevalence (with 95% CI) of phenotypic antimicrobial resistance against certain antibiotics within the BioProjects. Point and 95% CI estimates for major and very major error rates are presented in columns ME and VME. The dotted vertical lines represent the mean of the metrics within the given BioProject.

Antibiotics for which the NPV was estimable and reached 90% in at least half of the BioProjects (n=21): amikacin, azithromycin, ceftazidime, chloramphenicol, ciprofloxacin, colistin, ertapenem, fosfomycin, gentamicin, imipenem, kanamycin, meropenem, nalidixic acid, streptomycin, sulfamethoxazole, sulfisoxazole, temocillin, tetracycline, tigecycline, tobramycin, trimethoprim. Antibiotics for which the PPV was estimable and reached 90% in at least half of the BioProjects (n=15): ampicillin, cefepime, cefotaxime, cefoxitin, chloramphenicol, ciprofloxacin, colistin, fosfomycin, gentamicin, kanamycin, piperacillin, sulfamethoxazole, tetracycline, tobramycin, trimethoprim. Antibiotics for which the SE was estimable and reached 90% in at least half of the BioProjects (n=13): amikacin, ampicillin, ceftazidime, chloramphenicol, colistin, fosfomycin, gentamicin, kanamycin, streptomycin, sulfamethoxazole, tetracycline, tobramycin, trimethoprim. Antibiotics for which the SP was estimable and reached 90% in at least half of the BioProjects (n=29): amikacin, ampicillin, azithromycin, cefepime, cefotaxime, cefoxitin, cefpodoxime, ceftiofur, ceftriaxone, chloramphenicol, ciprofloxacin, colistin, ertapenem, fosfomycin, gentamicin, imipenem, kanamycin, levofloxacin, meropenem, nalidixic acid, nitrofurantoin, piperacillin, sulfamethoxazole, sulfisoxazole, temocillin, tetracycline, tigecycline, tobramycin, trimethoprim.

Antibiotics for which ME was estimable and did not exceed 5% in at least half of the BioProjects (n=28): ampicillin, azithromycin, cefepime, cefotaxime, cefoxitin, cefpodoxime, ceftiofur, ceftriaxone, chloramphenicol, ciprofloxacin, colistin, ertapenem, fosfomycin, gentamicin, imipenem, kanamycin, levofloxacin, meropenem, nalidixic acid, nitrofurantoin, piperacillin, sulfamethoxazole, sulfisoxazole, temocillin, tetracycline, tigecycline, tobramycin, trimethoprim. Antibiotics for which VME was estimable and did not exceed 5% in at least half of the BioProjects (n=15): amikacin, cefotaxime, ceftazidime, chloramphenicol, colistin, fosfomycin, gentamicin, kanamycin, nalidixic acid, nitrofurantoin, streptomycin, sulfamethoxazole, tetracycline, tobramycin, trimethoprim.

The ARG frequencies associated with the antibiotic identified in the metagenome analysis are summarised in Figure 3. Fig 4 shows the antibiotics for which no ARG was detected in the sample.

**Figure 3.**
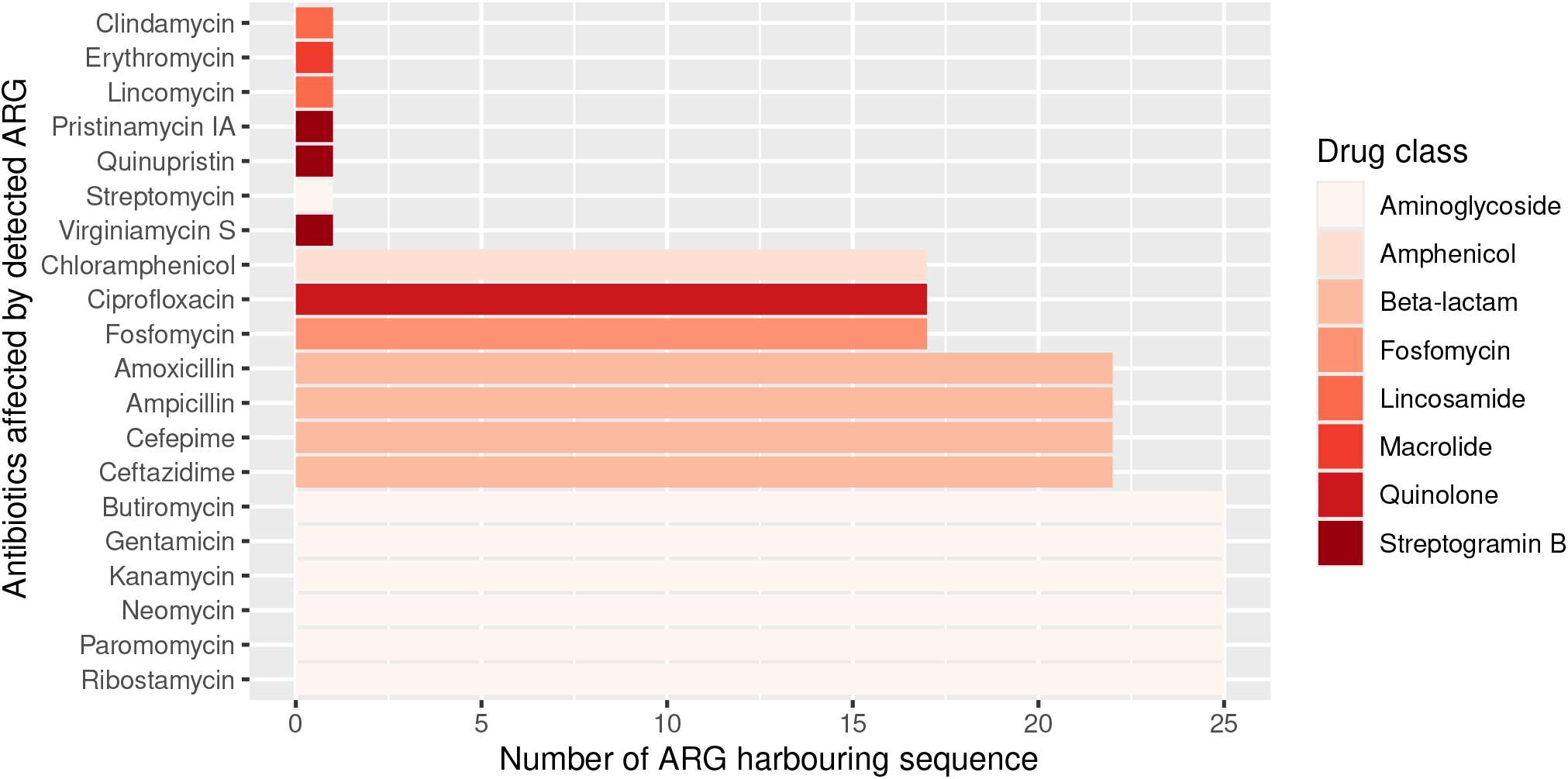
The number of sequences harboring ARGs by affected antibiotic compounds. The color of the bars represents the drug class to which a given antibiotic compound belongs.

**Figure 4.**
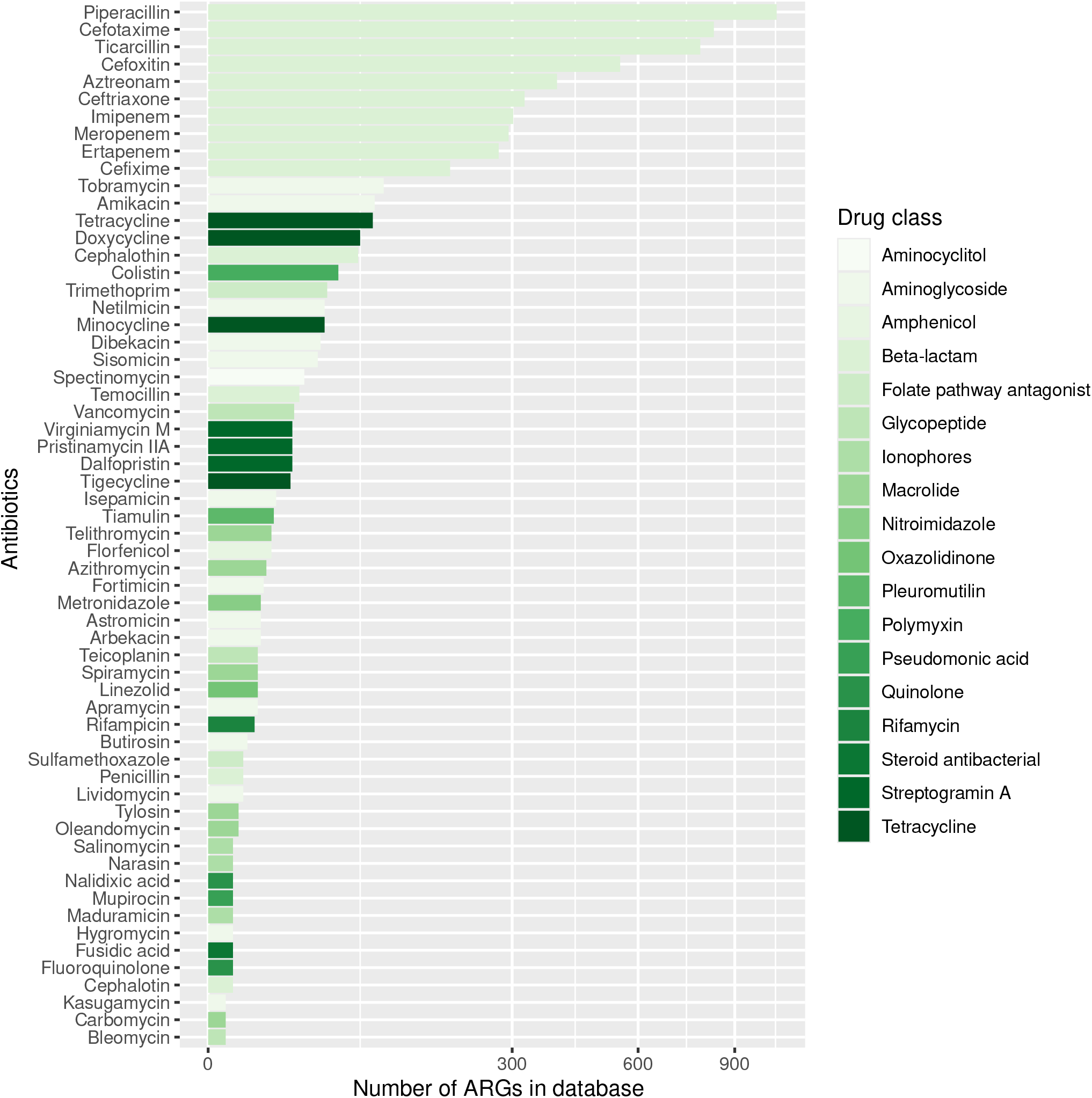
Antibiotics for which no ARG was detected in the sample are sorted by decreasing number of ARGs in the database affecting antibiotic compounds. The color of the bars represents the drug class to which a given antibiotic compound belongs. (cephalosporins, penams, penems, monobactams) that have very different properties from eachother or even within the subcategory (e.g. cephalosporin generations).^18^

## Discussion

Based on the phenotypic and genotypic resistance concordance analyses in *E. coli* strains, the following conclusions can be drawn. For 39% of antibiotics, sensitivity exceeded 90% in the evaluated BioProjects. In other words, for these antibiotics, phenotypically resistant strains are genotypically associated with a 90% probability of resistance. Based on the positive predictive values, a strain identified as genotypically resistant has at least a 90% probability of being phenotypically resistant for 45% of antibiotics. At 45% of antibiotics, there is less than a 5% probability that a phenotypically resistant strain will be identified as genotypically susceptible.

By 88% of antibiotics, phenotypically susceptible strains are at least 90% likely to be susceptible based on genotype. Strains detected as genotypically sensitive are at least 90% likely to be phenotypically sensitive at 64% of the antibiotics tested. The probability of identifying a phenotypically sensitive strain as resistant based on genotype is less than 5% for 85% of antibiotics.

To have more detailed understanding of the agreement results, a further aspect of challenges related to AST testing should be considered. Even broth microdilution, which is the AST method proposed by EUCAST (European Committee on Antimicrobial Susceptibility Testing) gives debatable results, and as do other classical, phenotype-based methods (e.g. disk diffusion). Dissimilarity in laboratory conditions, human resources, and discrepancies in laboratory techniques may lead to different results.^16^

A recent study including a machine learning-assisted ARG-based workflow for phenotypic AMR prediction indicates that the highest prediction accuracy for AMR against the drug classes tested appears by macrolides and sulfonamides. The highest uncertanity was found by beta-lactams that were also tested in combinations.^17^ From the demonstrated BioProjects, azithromycin, sulfamethoxazole and sulfisoxazole fall into the above categories. In contrast to the above mentioned publication, only sulfixazole yielded above average results considering all the investigated factors (NPV, PPV, SE, SP, ME, VME). In line with the above-mentioned study, rather large deviations from the mean were found by beta-lactams. However, this finding be explained by the large number of beta-lactams. In our case 14 different compounds were tested in the above-mentioned drug class (ampicillin, aztreonam, cefepime, cefotaxime, cefoxitin, cefpodoxime, ceftazidime, ceftriaxone, cefuroxime, ertapenem, imipenem, meropenem, piperacillin, temocillin). These antibiotics belong to separate subcategories within beta-lactams In contrast to single-strain cultures, similar comparisons using descriptive measures cannot theoretically be made with complex microbiological samples. One of the reasons for this is that only a small proportion of bacteria can be cultured and tested for phenotypic resistance.^6^ The recent advance of culturomics may partly address this aspect. Culturomics methods facilitates higher throughput, i.e. the identification of numerous difficult-to-culture bacteria, by applying several different growth conditions and media simultaneously.^19^ Even though, the number of condition combinations required for the same throughoutput can be experimentally reduced^**?**^, not all culturomics combinations are used in routine diagnostics. Thus, several bacteria involved in disease processes caused by a complex microbial communities can remain hidden, even if they play a critical role. However, shotgun metagenomic studies offer the opportunity to obtain data on the relative abundance rates of the taxa and, of particular clinical relevance, the genes, including ARGs ofcomplex microbiomes. The ARG prevalence data can then be used to gain insight into the AMRP of a given sample. Unfortunately, due to the limitations of bacteriological culturing, the concordance of AMRP and complex bacterial phenotypic resistance cannot currently be depicted. However, if we think of the complex microbiological samples as sets of bacterial single-strains, we can assume that the phenotypic and genotypic resistance/susceptibility matches for the strains may also be extended to them.

The fact that the study was carried out on *E. coli* strains may also be taken into account when evaluating the results. According to the publication by Hu and colleagues, in which phenotypic AMR profile prediction was performed using genotype-based machine learning algorithms by various bacterial species, predictions for *E. coli* strains were less robust and reliable compared to other species such as *Campylobacter jejuni* and *Enterococcus faecium*.^17^ These differences in phenotypic AMR profile prediction at the bacterial species level may also be relevant in bacterial communities containing multiple taxonomic groups. Furthermore, in addition to antimicrobial resistance at the cellular level, there is an increasing focus on the so-called cooperative resistance of whole bacterial communities. The dynamics within microbial communities (regardless of being single-species populations or mixed populations) can influence the response to antimicrobial therapy.^20^ A bacterial cell that can inactivate ABs with an exoenzyme can confer immunity to a cell that does not have the enzyme.^21^ Such interspecies interactions are already known, e.g. *Stenotrophomonas maltophilia* can also degrade imipenem in an enzyme-mediated manner, thereby protecting susceptible *Pseudomonas aeruginosa* in cystic fibrosis.^20^ Cooperative resistance is present in all microbe-rich environments.^21^ Most clinical samples are microbe-rich and therefore using genotypic resistance testing in practice should be considered for its beneficial, additional value.

From a clinical point of view, the question is which antibiotics are effective against the bacteria involved in a complex microbial community that induces or sustains a disease process. From the presented studies of *E. coli* strains, phenotypic susceptibility can be predicted with high confidence for the vast majority of antibiotics tested based on the ME, NPV, SP metrics of the genotype. If we extend these approach to metagenomic datasets, its clinical applicability for the selection of therapeutic agents could be considered. Reflecting to the AMRP based on the metagenomic analysis presented above, we can conclude the following. Even infections caused by complex microbial communities can be associated with antibiotics that were not affected by any ARGs found, despite being associated with ARGs in the reference database. The number of ARGs for different antibiotics and drug classes may vary widely in the different databases, and it is important to note that each ARG database is incomplete. However, we can assume that antibiotics against which no ARGs can be detected in a sample are more likely to be clinically effective if the ARG reference databases include a relatively higher number of ARGs against them.

Conventional phenotypic antimicrobial susceptibility testing procedures are still among the most significant methods for everyday routine bacteriological diagnostics. However, the adoption of nanopore sequencing, followed by comprehensive bioinformatic analysis, allows for rapid detection of microorganisms, along with the assessment of their abundance rates, presence of various genes, including ARGs or even virulence genes, possibly all within a timeframe of 4 to 24 hours.^22^ Consequently, clinical metagenomics based on Oxford Nanopore Technologies (ONT) has the potential to enhance diagnostic microbiology and clinical practices significantly. Nonetheless, the innovative nature of this approach raises several questions that have been addressed within this study, including discrepancies observed between the identified antimicrobial resistance genes and the expected phenotypic antimicrobial resistance.^23^

## Declarations

### Ethics approval and consent to participate

Not applicable.

### Consent for publication

Not applicable.

### Data Availability

The raw long-read data are available from the corresponding author upon reasonable request.

### Author contributions statement

NS takes responsibility for the data’s integrity and the data analysis’s accuracy. AGT and NS conceived the concept of the study. AGT and TN collected the biological samples. AGT and NS participated in the bioinformatic analysis. AGT, NS and SÁN participated in the drafting of the manuscript. AGT, CF, IC, NS, SÁN, TN, and TR critically revised the manuscript for important intellectual content. All authors read and approved the final manuscript.

## Authors’ information

Not provided.

## References

1. Muteeb, G., Rehman, M. T., Shahwan, M. & Aatif, M. Origin of antibiotics and antibiotic resistance, and their impacts on drug development: A narrative review. Pharmaceuticals 16, 1615 (2023).

2. Magnusson, U. Prudent and effective antimicrobial use in a diverse livestock and consumer’s world. J. Animal Sci. 98, S4–S8 (2020).

3. Motro, Y. & Moran-Gilad, J. Next-generation sequencing applications in clinical bacteriology. Biomol. Detect. Quantification 14, 1–6 (2017).

4. Khan, Z. A., Siddiqui, M. F. & Park, S. Current and emerging methods of antibiotic susceptibility testing. Diagnostics 9, 49 (2019).

5. Boolchandani, M., D’Souza, A. W. & Dantas, G. Sequencing-based methods and resources to study antimicrobial resistance. Nat. Rev. Genet. 20, 356–370 (2019).

6. Steen, A. D. et al. High proportions of bacteria and archaea across most biomes remain uncultured. The ISME journal 13, 3126–3130 (2019).

7. Stewart, E. J. Growing unculturable bacteria. J. Bacteriol. 194, 4151–4160 (2012).

8. Van Belkum, A. et al. Rapid clinical bacteriology and its future impact. Annals Lab. Medicine 33, 14 (2013).

9. Chibucos, M. C. et al. An ontology for microbial phenotypes. BMC Microbiol. 14, 1–8 (2014).

10. Pataki, B. Á. et al. Understanding and predicting ciprofloxacin minimum inhibitory concentration in Escherichia coli with machine learning. Sci. Reports 10, 15026 (2020).

11. Li, D., Liu, C.-M., Luo, R., Sadakane, K. & Lam, T.-W. Megahit: an ultra-fast single-node solution for large and complex metagenomics assembly via succinct de bruijn graph. Bioinformatics 31, 1674–1676 (2015).

12. De Coster, W., D’hert, S., Schultz, D. T., Cruts, M. & Van Broeckhoven, C. NanoPack: visualizing and processing long-read sequencing data. Bioinformatics 34, 2666–2669 (2018).

13. Bortolaia, V. et al. Resfinder 4.0 for predictions of phenotypes from genotypes. J. Antimicrob. Chemother. 75, 3491–3500 (2020).

14. FDA, US. Class II special controls guidance document: antimicrobial susceptibility test (AST) systems. Rockville, MD: US FDA (2009).

15. International Organization for Standardization. Clinical Laboratory Testing and in Vitro Diagnostic Test Systems-Susceptibility Testing of Infectious Agents and Evaluation of Performance of Antimicrobial Susceptibility Test Devices: Reference Method for Testing the in Vitro Activity of Antimicrobial Agents Against Rapidly Growing Aerobic Bacteria Involved in Infectious Diseases (ISO, 2006).

16. Rebelo, A. R. et al. One day in Denmark: comparison of phenotypic and genotypic antimicrobial susceptibility testing in bacterial isolates from clinical settings. Front. Microbiol. 13, 804627 (2022).

17. Hu, K. et al. Assessing computational predictions of antimicrobial resistance phenotypes from microbial genomes. Briefings Bioinforma. 25, bbae206 (2024).

18. Bush, K. & Bradford, P. A. β -lactams and β -lactamase inhibitors: an overview. Cold Spring Harb. Perspectives Medicine 6, a025247 (2016).

19. Lagier, J.-C. et al. The rebirth of culture in microbiology through the example of culturomics to study human gut microbiota. Clin. Microbiol. Rev. 28, 237–264 (2015).

20. Denk-Lobnig, M. & Wood, K. B. Antibiotic resistance in bacterial communities. Curr. Opin. Microbiol. 74, 102306 (2023).

21. Vega, N. M. & Gore, J. Collective antibiotic resistance: mechanisms and implications. Curr. Opin. Microbiol. 21, 28–34 (2014).

22. Ring, N. et al. Rapid metagenomic sequencing for diagnosis and antimicrobial sensitivity prediction of canine bacterial infections. Microb. Genomics 9, 001066 (2023).

23. Yee, R., Dien Bard, J. & Simner, P. J. The genotype-to-phenotype dilemma: how should laboratories approach discordant susceptibility results? J. Clin. Microbiol. 59, 10–1128 (2021).

